## Supplementary figures and images for "Myofibroblast Senescence Promotes Arrhythmogenic Remodeling in the Aged Infarcted Rabbit Heart"

### Supplemental Figure 1

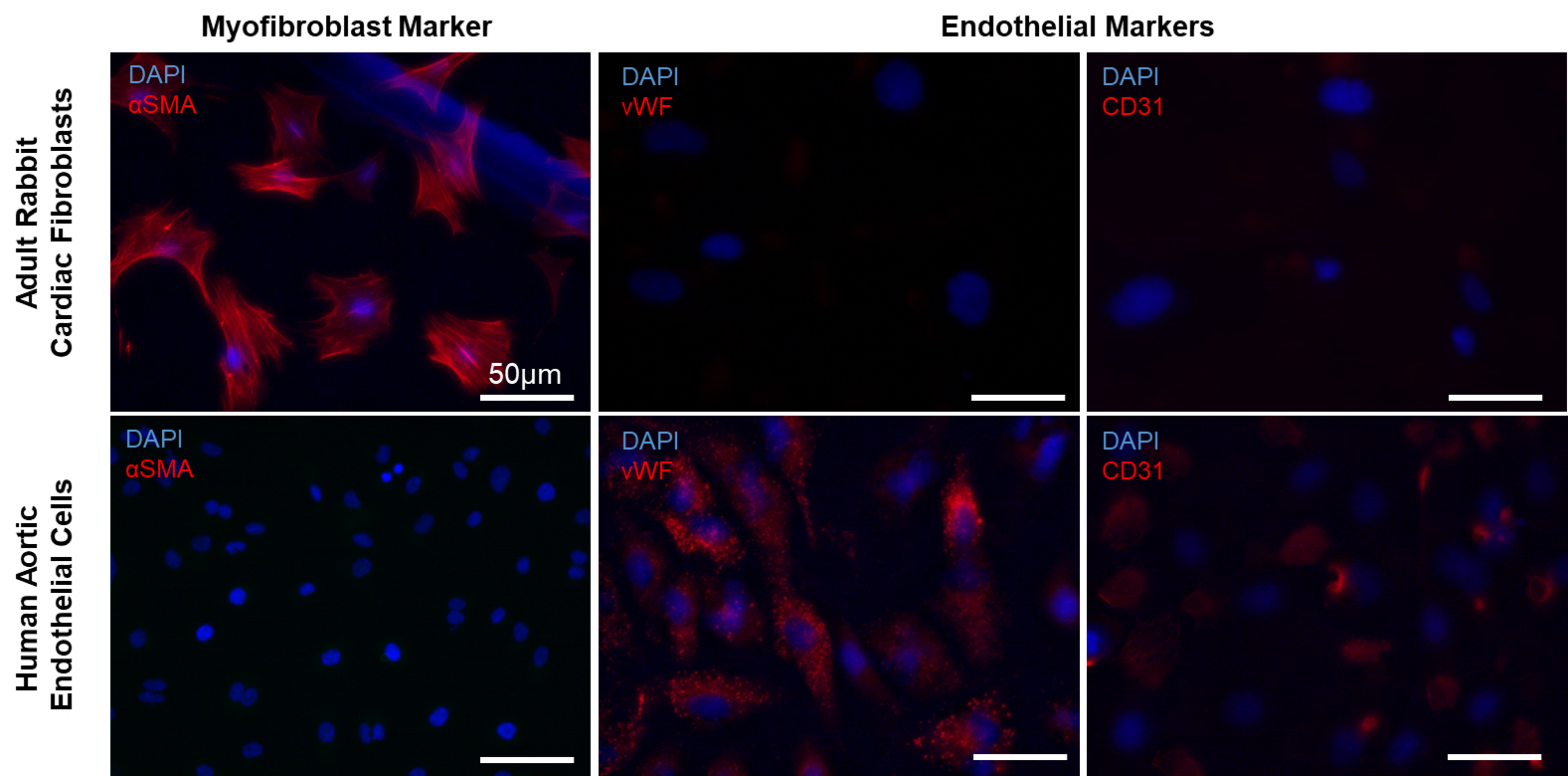
