## Supplemental Figure 2 for "Myofibroblast Senescence Promotes Arrhythmogenic Remodeling in the Aged Infarcted Rabbit Heart"

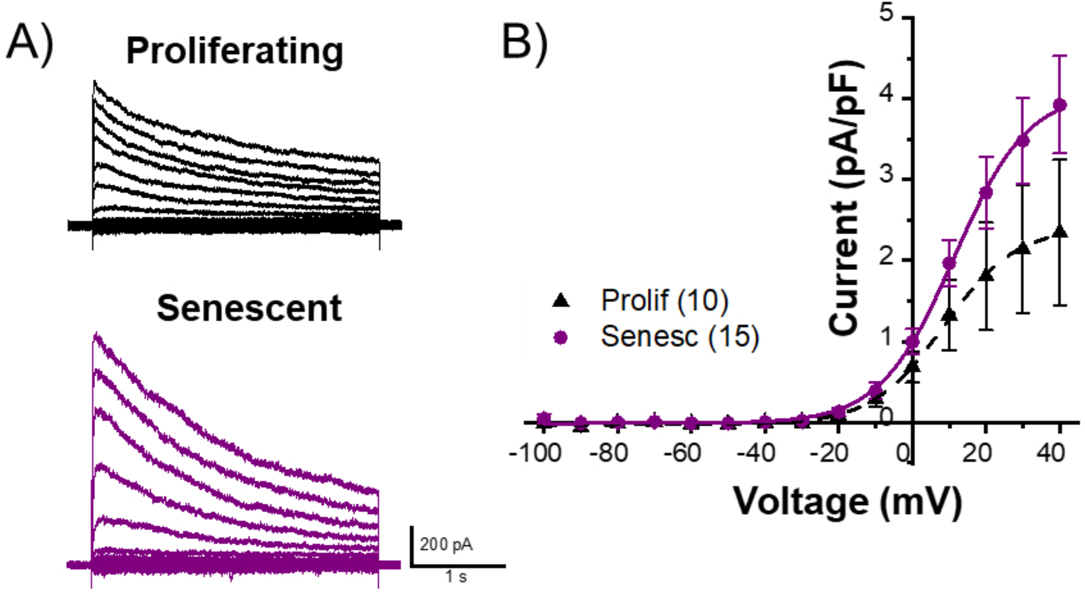


C)

$$I_{Fib}(V) = A_2 + \frac{A_1 - A_2}{1 + e^{(V - V_0)/dV}}$$

$$I_{Pro}(V) = gPro * \left(2.4 + \frac{-0.03 - 2.4}{1 + e^{(V - 8.5)/10}}\right)$$

$$I_{Sen}(V) = gSen * \left(4.1 + \frac{-0.01 - 4.1}{1 + e^{(V - 11.3)/10}}\right)$$

$$I_{Pro}(V)_{Shift} = gPro * \left(2.4 + \frac{-0.03 - 2.4}{1 + e^{(V - 27.5)/10}}\right)$$

$$I_{Sen}(V)_{Shift} = gSen * \left(4.1 + \frac{-0.01 - 4.1}{1 + e^{(V - 37.5)/10}}\right)$$
