## Supplemental Table 1 for "Myofibroblast Senescence Promotes Arrhythmogenic Remodeling in the Aged Infarcted Rabbit Heart"

| Target | Host Species | Company | Product Number |
|---|---|---|---|
| αSMA | Mouse | Sigma | A5228-100UL |
| Troponin T | Mouse | Invitrogen | Ma5-12960 |
| Cx43 | Goat | Abcam | ab219493 |
| γH2AX | Rabbit | Cell Signaling | 2577 |
| γH2AX | Mouse | Abcam | ab26350 |
| αSMA | Goat | Novus | NB300-978SS |
| CD31 | Goat | R&D Systems | AF3628 |
| vWF | Sheep | Abcam | ab11713 |
| WGA Alexa 594 | | Thermo | W11262 |
| Mouse Alexa Fluor 647 | Goat | Thermo | A21235 |
| Goat Alexa Fluor 488 | Donkey | Thermo | A11005 |
| Rabbit Alexa Fluor 594 | Goat | Thermo | A110012 |
| Sheep Alexa Fluor 594 | Donkey | Thermo | A110016 |
| Sheep Alexa Fluor 647 | Donkey | Thermo | A21448 |
