## Supplemental Table 2 for "Myofibroblast Senescence Promotes Arrhythmogenic Remodeling in the Aged Infarcted Rabbit Heart"

| Target | Primer/probe Sequences (5'-3') | Efficiency |
|---|---|---|
| SRP14 | Accession: XM_002717984.2<br>Forward: 5' – TTCCAGAAATGCCGGTTGTC – 3'<br>Reverse: 5' – CAAAGCCCTCCACAGAACCTT – 3' | 102% |
| p16 | Accession: XM_008255019.2<br>Forward: 5' – GCCGAAGGCGGTAACCA -3'<br>Reverse: 5' – GCTTCGATGCTGGACTTCTCA – 3' | 96% |
| p21 | Accession: XM_002714669.2<br>Forward: 5' – GAGACTGCGACGCACTCATG – 3"<br>Reverse: 5' – CGCTCCCAGGCGAAGTT – 3" | 98% |
| p53 | Accession: XM_017348089.1<br>Forward: 5' – CCTGGAGCTCTGGTCTGAGATC – 3'<br>Reverse: 5' – CCACCACGTCAAATACCCCTAA – 3' | 100% |
| TGFβ | Accession: XM_008249704.2<br>Forward: 5' – CACCCGCGTGCTAATGGT – 3'<br>Reverse: 5' – CCGGAGCTCTGACGTGTTAAA – 3' | 97% |
| IL-6 | Accession: NM_001082064.2<br>Forward: 5' – GAAAACACCAGGGTCAGCAT – 3'<br>Reverse: 5' – TGAGGGTGGCTTCTTCATTC – 3' | 107% |
| CCL2 | Accession: NM 001082294.1<br>Forward: 5' – TGAATTCCCCAGTCACCTGC – 3'<br>Reverse: 5' – TTGGGACACTTGGTGCTGTT – 3' | 97% |
| CXCL2 | Accession: XM_002717021.3<br>Forward: 5' – AGACCGTGCAGGGCATTC – 3'<br>Reverse: 5' – GACTTGAGCGTGGCTATGACTTC – 3' | 101% |
| MMP9 | Accession: NM_001082203.1<br>Forward: 5' – TCCAACTTTGACAGCGACA – 3'<br>Reverse: 5' - GGAGTGATCCAAGCCCAGTG – 3' | 109% |
| TIMP1 | Accession: XM_008272340.2<br>Forward: 5' – GCAACTCCGACCTTGTCATC – 3'<br>Reverse: 5' – AGCGTAGGTCTTGGTGAAGC – 3' | 100% |
| TIMP2 | Accession: XM_008252510.2<br>Forward: 5' – GCGAGAAGGAGGTGGACT – 3'<br>Reverse: 5' – TTGTCGGGGCCTTTGAACAT – 3' | 103% |
| IL-8 | Accession: NM_001082293.1<br>Forward: 5' –CTGTTGGTCAGGCCATGAGT – 3'<br>Reverse: 5' –AAAGTGCTTCCATGTGCCCT – 3' | 104% |
| SDHA | Accession: XM_002723194<br>Forward: 5' –GGACCAGGACGCCATCCACTAC – 3'<br>Reverse: 5' –TCCACCGAACGCACGCTGATAG – 3' | 106% |
| HPRT1 | Accession: NM_001105671<br>Forward: 5' –CCTTGGTCAAGCAGTATAATC – 3'<br>Reverse: 5' –GGGCATATCCTACAACAAAC – 3' | 101% |
